# The role and origins of human attitudes in Human–Wildlife Conflict responses: Insights from Spectacled Bears (*Tremarctos ornatus*) and other wild carnivores in Southern Ecuador

**DOI:** 10.64898/2026.03.29.715142

**Authors:** Francisco Lopes, Mateo Peñaherrera-Aguirre, Rodrigo Cisneros

## Abstract

**Background:** Human-wildlife conflict, which motivates retaliatory killings, is a major driver of species decline globally. Addressing an open question in human–wildlife conflict, we test whether evolutionary-rooted human attitudes, independent of economic losses, better predict retaliatory responses.

**Methods:** We examined human attitudes toward spectacled bears (*Tremarctos ornatus*) and other wild carnivores in a wildlife conflict-zone in southern Ecuador by conducting interviews in rural communities. We measured both established variables - such as education levels, age, and gender - and novel psychometric variables to identify predictors of human-wildlife conflict responses.

**Results:** Perceptions of animals emerged as the strongest predictor of conflict responses. Communities exhibiting high levels of vengefulness, particularly within an animal-directed Culture of Honor, where individuals, especially men, are expected to respond strongly or violently to perceived threats, were more likely to support lethal interventions. Conversely, individuals with strong environmental education backgrounds demonstrated more positive perceptions of wildlife, highlighting education’s potential role in conflict mitigation.

**Conclusion:** Evolutionary-derived attitudes, rather than economic factors, primarily drive human responses to wildlife conflict. Effective strategies to reduce violence against wildlife should incorporate human perceptions and culturally rooted values to address the underlying social and psychological drivers of conflict.

## Introduction

Interactions with wildlife have been a constant feature of human existence. Competition for habitat and resources with other wildlife played a crucial role in enabling *Homo sapiens* to emerge as the most dominant ecological force on the planet (Waters et al., 2016) resulting in human relationships with wildlife deeply embedded in the evolutionary history of our species. Although some interactions are considered beneficial (Nyhus, 2016), such as the domestication of animals, a substantial proportion of human-wildlife interactions have evolved into Human-Wildlife Conflict (HWC), that is, situations where the needs, behaviors, or activities of humans and wildlife come into direct opposition, resulting in negative consequences for one or both (Conover, 2002). HWC is now recognized as one of the leading drivers of species decline and population extirpation (Gross et al., 2021).

HWC can have profound implications for humans, adversely affecting human life, safety, economic security, food supply, health, or property (Treves & Karanth, 2003; Peterson et al., 2011). These outcomes often result from economic losses incurred due to damage inflicted on crops, livestock, cinegenic species, and property (Nyhus, 2016). This impact is disproportionately felt in rural communities, which frequently depend almost entirely on agricultural and pastoral livelihoods, and may suffer irreparable economic setbacks as a result (Cisneros et al., 2025; Braczkowski et al., 2023).

HWC also affects wildlife, which frequently faces intense persecution in response to conflict, leading to the widespread killing of animals (Nyhus, 2016). Such effects are particularly evident in the case of large carnivores and herbivores that are often subject to the highest levels of hostility due to both real and perceived threats to human safety and livelihoods (Chapron et al., 2014; Saberwal et al., 1994). The targeting of these species not only reduces their populations but also fragments their habitats and restricts their ranges (Woodroffe et al., 2005). This is especially concerning because large-bodied mammals typically occur at low population densities, require extensive home ranges, and are acutely vulnerable to habitat degradation and fragmentation (Robinson & Redford, 1986; Noss et al., 1996). These losses can also trigger ecological cascades with far-reaching consequences (Strong & Frank, 2010) and amplify the risk of ecosystem-level disruptions, as illustrated by the extirpation of wolves (*Canis lupus*) in Yellowstone National Park, which produced catastrophic effects (Ripple & Beschta, 2011). HWC in the Andean region of Ecuador poses a serious and escalating threat to carnivore populations, with the spectacled bear (*Tremarctos ornatus*) and other large predators often disproportionately targeted due to fear, misperceptions, and retaliatory killings. Within the present study area, HWC is a significant and escalating concern. Conflict has been recorded across a diverse range of species, including large and small carnivores, raptors, and wild herbivores. Nevertheless, carnivores appear to experience the most severe consequences (Cisneros et al., 2025; MAE, 2019; Vela-Vargas et al., 2021; García-Rangel, 2012). The spectacled bear (*Tremarctos ornatus*) is undergoing marked population declines, and in the Ecuadorian Andes, specialists cite retaliatory killings linked to HWC as a primary driver of this trend (Tirira, 2021; MAE, 2019; García-Rangel, 2012). The species is currently listed as Endangered (EN) in the most recent Ecuadorian Red List of Mammals (Tirira, 2021), reflecting the gravity of the situation. Moreover, the spectacled bear is not alone in facing these pressures. Other carnivores in the region, such as the puma (*Puma concolor*), the Andean fox (*Lycalopex culpaeus*), and the jaguar (*Panthera onca*), are also experiencing serious conservation threats and are included among the endangered species of the Ecuadorian Andes (Tirira, 2021).

In the Andes, including our study area, local communities often blame the spectacled bear for cattle deaths, yet these losses are frequently exaggerated (Goldstein et al., 2006). Growing evidence shows that fear and misperception - rather than verified crop or livestock damage - drive many bear killings: in rural Bolivia, bears were labeled agricultural pests despite no confirmed presence or losses, while most crop damage was caused by parrots, parakeets, and skunks (Albarracín & Aliaga-Rossel, 2018). A regional review likewise found bears routinely blamed for livestock deaths and disappearances, prompting poaching unlinked to verified conflict (Goldstein et al., 2006). Reports from Peru and Bolivia document rising bear mortality - including females with cubs and juveniles - killed without any suspicion of damage; in some cases, carcasses were shown to tourists for a fee (Figueroa, 2008; Albarracín, 2010; Albarracín et al., 2013).

However, retaliation explains only part of wildlife killings: many bear deaths occur without verified damage or subsistence need, driven by fear, folklore, and symbolism (Goldstein et al., 2006), as happens for many other wildlife. Examples include Peru’s Yawar Fiesta condor–bull rite (Barnes, 1994), poaching of barn owls in Costa Rica (Enríquez & Mikkola, 1997), giant anteaters in Brazil (Catapani et al., 2023), snakes in Bahia (Fita et al., 2010), and bats near Paraíba (Rego et al., 2015). In the Ecuadorian Andes, particularly in the Cayambe region, disturbing events have been documented, such as the crucifixion of a spectacled bear (Moscoso, 2020). In all of these cases, extreme and apparently unnecessary violence is used, indicating motivations for killing beyond economic retaliation. These patterns indicate that attitudes and perceptions, not just economic loss, shape HWC. Consistent with this, perceived risk and fear strongly predict carnivore attitudes beyond documented losses (Bhatia et al., 2020). Thus, individual differences, including emotions, psychometrics, and evolutionary influences, should be examined alongside age, environmental education, and religion (Baldry, 2003; Gullone & Robertson, 2008; Chikezie et al., 2023; Pasaribu et al., 2021; White, 1967).

The Animality framework can capture thoughts, feelings, and behaviors toward animals, distinct from general personality, and comprising Emotional Regard and Attraction to Animals, together forming a General Factor of Animality that predicts empathy across taxa (Steklis et al., 2018, 2023; Figueredo et al., 2023; Peñaherrera-Aguirre, 2024; Gunnarsson & Hedblom, 2023). Evidence shows perceptions often disproportionate to conflicts: prairie dogs and Zanzibar red colobus are vilified despite benefits to production systems; small species cause greater crop loss than large, visible ones; historic events and media amplify fear (Reading et al., 2009; Siex & Struhsaker, 1999; Naughton-Treves & Treves, 2005; Dickman, 2010; Knight, 2000; Linnell et al., 2003; Prokop et al., 2009). Culture of Honor further predicts aggressive responses to threats to property or reputation, especially in pastoral settings, potentially fueling indiscriminate, illegal, and ecologically harmful retaliation (Nisbett & Cohen, 1996; Figueredo et al., 2004; Shackelford, 2005; Nyhus, 2016), as seen with retributive leopard killings in South Africa (Nyhus, 2016). In the present study we test how established factors (age, environmental education, religion) and proposed drivers (perceptions, emotions, General Factor of Animality, and Culture of Honor) jointly shape attitudes toward wildlife and responses to HWC.

We address the gap in HWC studies that focus on economic motivations for wildlife retaliatory killing by examining the largely un-explored role of evolutionary-rooted human attitudes. We test the hypothesis that perceptions and culturally embedded values better predict support for retaliation than material impacts. Focusing on spectacled bears and other carnivores in rural southern Ecuador, we conducted interviews and measured established factors (economic impact, environmental education, age, gender) alongside novel psychometrics - the General Factor of Animality and Culture of Honor - analyzed with a Sequential Canonical Model. The approach used enabled examining conflict mitigation through the lens of human psychology and culture, not economics alone.

## Materials & Methods

### Participants and Measures

The study was conducted in various populated centers within the provinces of Zamora Chinchipe, Loja, and El Oro, located in the southern region of Ecuador. The study began with an exploratory phase to gather information regarding reported HWC, due to the lack of previous research in the area. Once affected individuals were identified, in situ visits were conducted, employing a chain sampling strategy (Newing, 2010) to carry out semi-structured interviews with farmers and individuals recommended by them. Local farmers from the communities were interviewed to investigate all aspects outlined in the study’s objectives. The sample included individuals who had experienced HWC as well as neighbors from the same area who had not experienced such conflicts in the last five years, ensuring a balanced representation of perspectives from both groups.

We collected questionnaire data from 152 participants living near wildlife reserves in Southern Ecuador. In addition to sociodemographic information (sex, age, participants’ education, parental education), data were collected on December 2024 through March 2025 using the following questionnaires:

The Environmental Education Scale (EES) evaluates the extent of environmental education an individual has received throughout their life, covering broad thematic areas such as care for the environment and forests, as well as care for nature and wildlife. The EES consists of eight items, allowing for the measurement of an individual’s level of environmental education and facilitating comparisons across participants. Each item was rated on a 5-point Likert scale, with higher scores indicating greater exposure to and engagement with environmental education. The measurement instrument used for this variable was developed by the authors specifically for the present study.

The Centrality of Religiosity Scale (CRS-10; Huber & Huber, 2012) evaluates the centrality of religion in an individual’s life across five core dimensions: intellectual interest, ideological beliefs, public practice, private practice, and religious experience. The CRS-10 consists of ten items, allowing for a nuanced measurement of religious involvement and significance.

Participants responded to each item using a 5-point Likert scale, with higher scores indicating greater religiosity. The Centrality of Religiosity Scale (CRS-10; Huber & Huber, 2012) is distributed under a Creative Commons Attribution 4.0 International License; accordingly, it was used in this study in full compliance with its licensing terms.

The Revenge Subscale of the Culture of Honor Questionnaire (Figueredo et al., 2004) was used to measure participants’ association with Culture of Honor. The original instrument included 32 social vignettes requiring participants to indicate whether the behavior of the protagonist featured in each vignette should be considered an underreaction or an overreaction. It uses a 6-point Likert scale. We adapted the scale for its cross-cultural use (human-directed Culture of Honor).

Similarly, the present study transformed the latter items wherein the target of the offense or harm was no longer a person but an animal (animal-directed Culture of Honor). In these adapted versions, lower scores indicate an underreaction and, consequently, a greater Culture of Honor, whereas higher scores indicate an overreaction, thus a lower Culture of Honor. The authors have permission to use this instrument from the copyright holders (Figueredo et al., 2004).

The Animality Questionnaire was used to measure Animality, a construct that encompasses variations in individuals’ thoughts, feelings, and behaviors towards animals and is distinct from human personality traits (Peñaherrera-Aguirre et al., 2024). Animality is related to the concept of biophilia, which suggests that humans have an innate affinity for bonding with nature (Gunnarsson & Hedblom, 2023). This instrument includes 30 items. The version used in this study employed a 5-point Likert scale. The authors have permission to use this instrument from the copyright holders (Steklis et al., 2018).

The present study also asked participants to report on their perceptions of native animals - opossums (*Didelphis spp*.), spectacled bears (*Tremarctos ornatus*), Andean foxes (*Lycalopex culpaeus*), pumas (*Puma concolor*), tayras (*Eira barbara*), weasels (*Mustela frenata*), feral dogs (*Canis lupus familiaris*), and hawks (*Geranoaetus melanoleucus*) - based on their degree of agreement with a list of 15 items describing each species. The instrument used a 5-point Likert scale. The measurement instrument used for this variable was developed by the authors specifically for the present study.

Participants were also asked to report the amount of hate they felt towards each native animal (opossums, spectacled bears, Andean foxes, pumas, tayras, weasels, feral dogs, and hawks) using a 5-point Likert scale. The measurement instrument used for this variable was developed by the authors specifically for the present study.

The Responses to Wildlife scale was measured by presenting participants with a list of seven potential behavioral reactions considering a hypothetical scenario in which the individual encounters a particular animal in the wild. Responses ranged from fleeing (lowest score) the area to hunting the animal (highest score). The species featured in these scenarios included opossums, spectacled bears, Andean foxes, pumas, tayras, weasels, feral dogs, and hawks. The measurement instrument used for this variable was developed by the authors specifically for the present study.

The ethical committee of the University of Porto approved this study. (Ref.: Proc. CE2025/p200). Written informed consent was obtained from all participants prior to data collection.

### Statistical Analysis

All items were standardized prior to examining each scale’s psychometric proprieties. Internal consistency values were examined with Cronbach’s α, and the corresponding factor structure was established following unit-weighted factor estimations with factor loadings computed as part-whole correlations between the standardized unit-weighted factor scores and its standardized items. All psychometric examinations were conducted in SAS 9.4 (SAS Institute Inc., 2015).

These standardized factor scores were later exported to UniMult 2.0 (Gorsuch, 2016) and were subsequently included as part of a Sequential Canonical Cascade model. This model procedure is established as a sequence of equations organized as a series of steps, wherein a variable operating as a criterion becomes a predictor in the following step of the analysis. The model employs a hierarchical partitioning of variance (Type 1 Sum of Squares) based on multiple F-tests, reducing the risk of Type I errors (Figueredo et al., 2017).

## Results

### Descriptive statistics

Of the 149 participants who participated in the study, 64 were females, and 85 were males. The sample mean age was 48.8 years (+ 14.2 years SD). Most participants either completed primary education (*n* = 49), obtained a high school diploma (*n* = 57), or held a bachelor’s degree (*n* = 28); a smaller number of participants either held a postgraduate degree (*n* = 8) or had no formal education (*n* = 7).

### Psychometric Validation

The religiosity scale exhibited a high level of internal consistency (α = 0.922) alongside adequate factor loadings, indicating a satisfactory level of structural validity (Figure 1a). The environmental education dimension exhibits high levels of internal consistency (α = 0.947), with adequate factor loadings (Figure 1b). The human-directed Culture of Honor scale exhibited a high level of internal consistency (α = 0.949) and was accompanied by adequate factor loadings (Figure 2a). The animal-directed Culture of Honor also exhibited a high level of internal consistency (α = 0.958) and was accompanied by adequate factor loadings (Figure 2b). The psychometric models revealed that the emotional regard dimension loaded positively and significantly onto the “enjoys-loves” (r = 0.948, p <.0001) and “affectionate” (r = 0.940, p <.0001) parcels. Similarly, the attraction to animal’s dimension loaded positively and significantly onto the intellectual curiosity (r = 0.884, p <.0001), special connection (r = 0.877, p <.0001), and understanding (r = 0.871, p <.0001) parcels. Additionally, the analysis supported the presence of a General Factor of Animality, which loaded positively and significantly onto the emotional regard and attraction to animals’ dimensions (r = 0.948, p <.0001).

**Figure 1.**
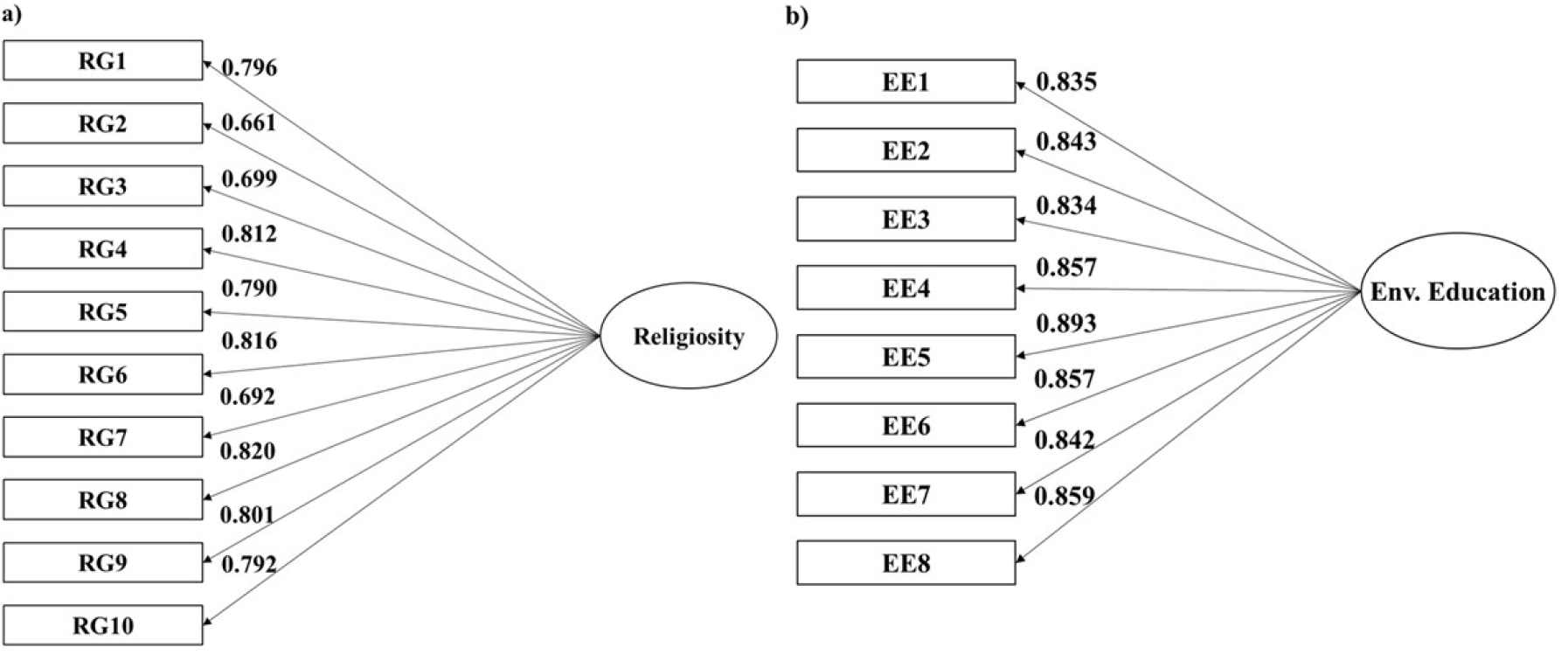
Latent structure of a) Religiosity; and b) Environmental education.

**Figure 2.**
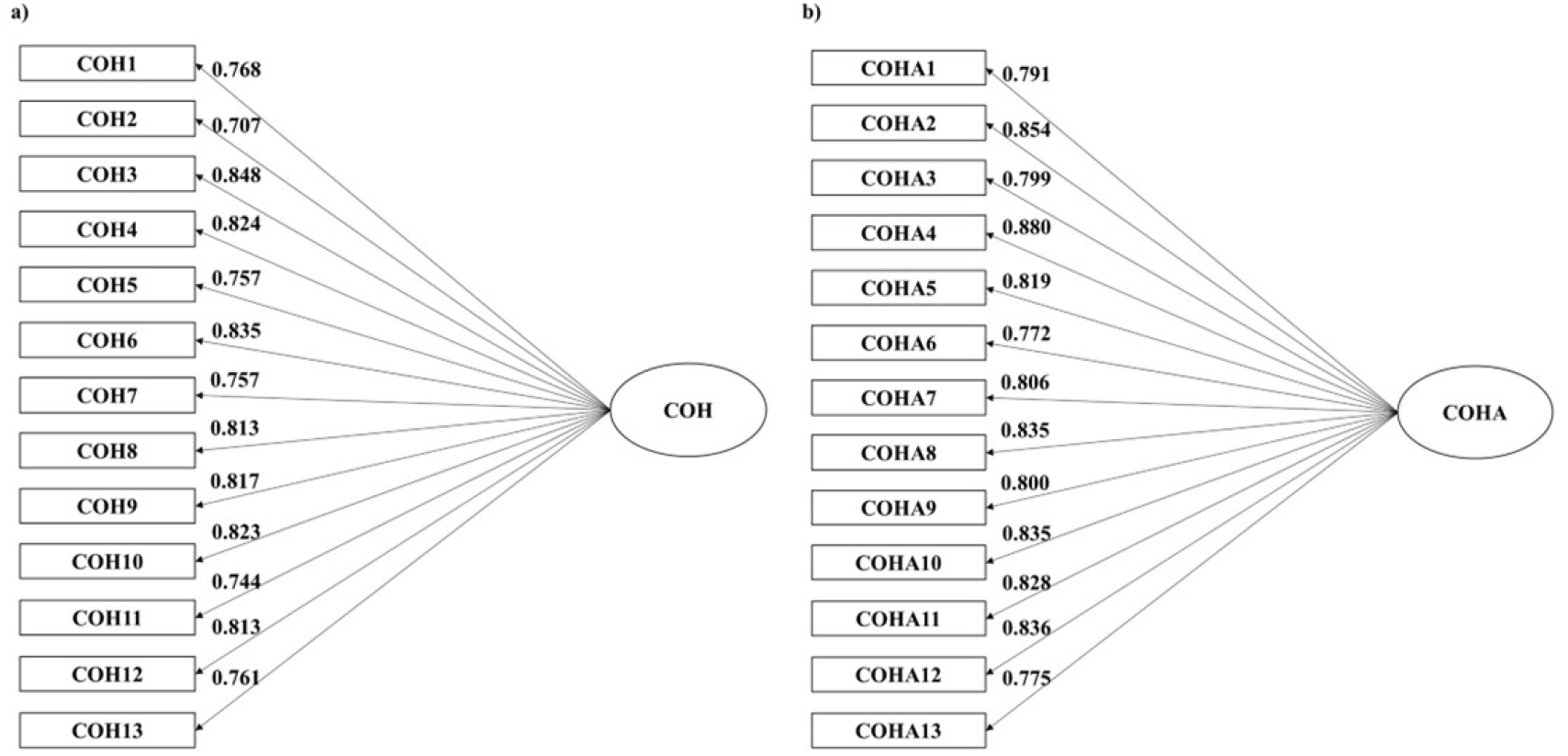
Latent structure of a) Human-directed culture of honor; and b) Animal-directed culture of honor unit-weighted factor.

A general dimension of negative perceptions towards animals also exhibited a high level of internal consistency (α = 0.909), with this latent structure loading positively and substantially onto all indicators (Figure 3). The responses to the conflict factor displayed a high level of internal consistency (α = 0.804), with adequate factor loadings (Figure 4). Overall, these results strongly suggest that all the instruments used in this study had strong psychometric properties. Consequently, these variables were included in a subsequent structural model to determine their influence on participants’ responses to conflict.

**Figure 3.**
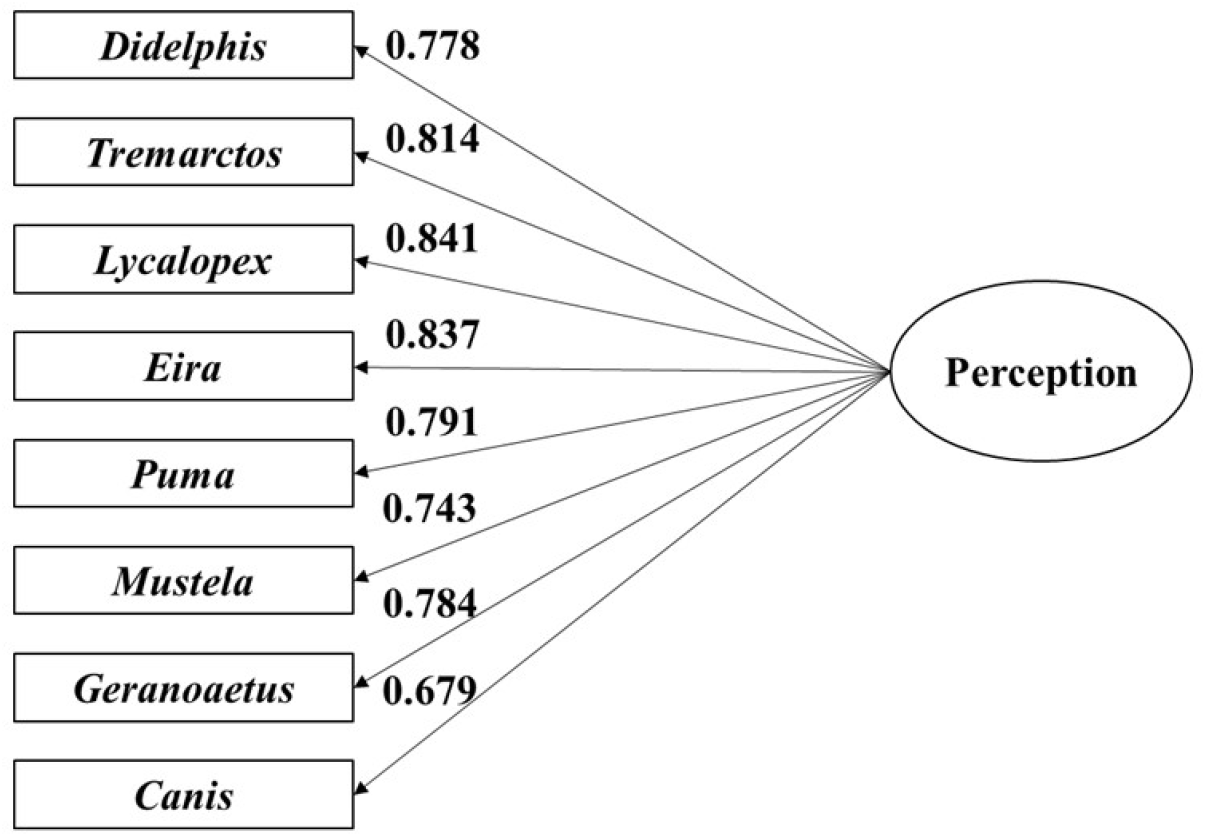
Latent structure of an animal perception unit-weighted factor.

**Figure 4.**
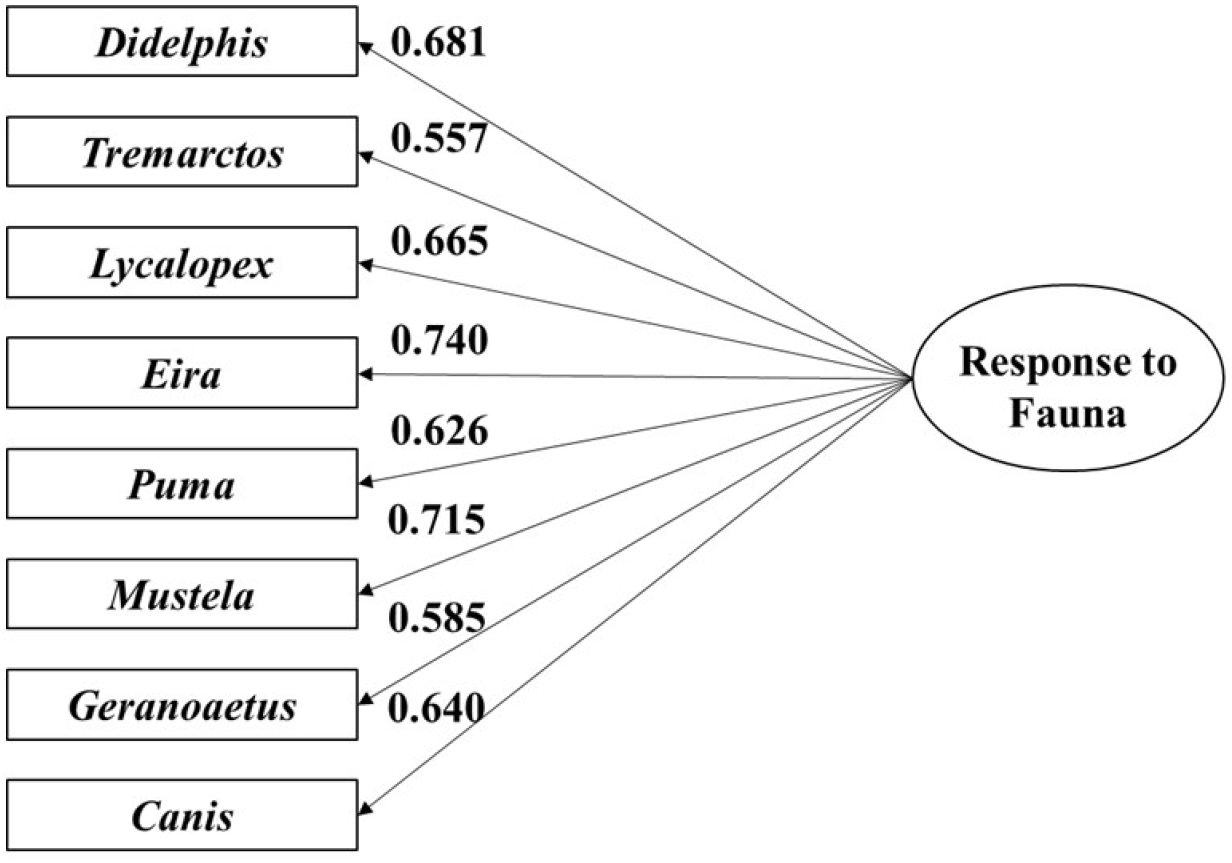
Latent structure of a response to fauna unit-weighted factor.

### Structural path results

The overall Sequential Canonical Cascade Analysis reached statistical significance (p<.0001) and explained 33% of the variance. A post-hoc power analysis conducted using the statistical platform GPower 3.1 (Erdfelder et al., 2009) indicated that the current sample achieved an adequate level of statistical power (1-β = 0.907).

Table 1 and Figure 5 describe these results in more detail. The Culture of Honor scale employs a Likert scale that ranks participants’ levels of agreement, with lower scores indicating that the protagonist’s behavior underreacted to the social scenario, whereas higher scores suggest that the protagonist overreacted. Hence, the human-directed Culture of Honor was positively predicted by male gender, religiosity, and education.

**Table 1.**
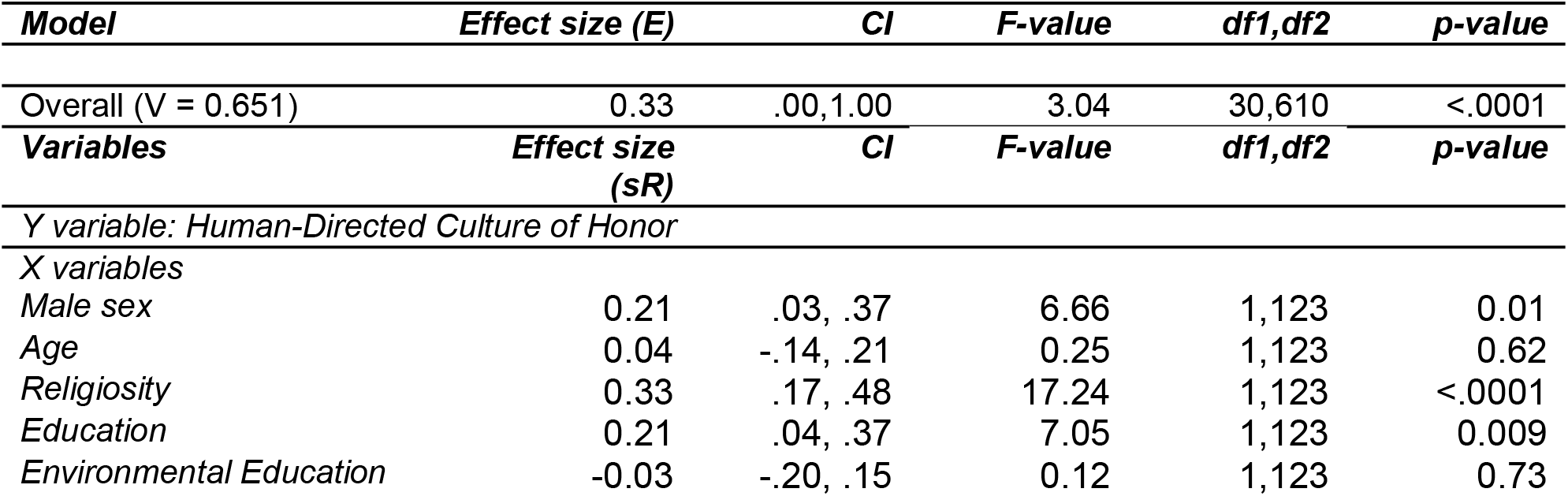
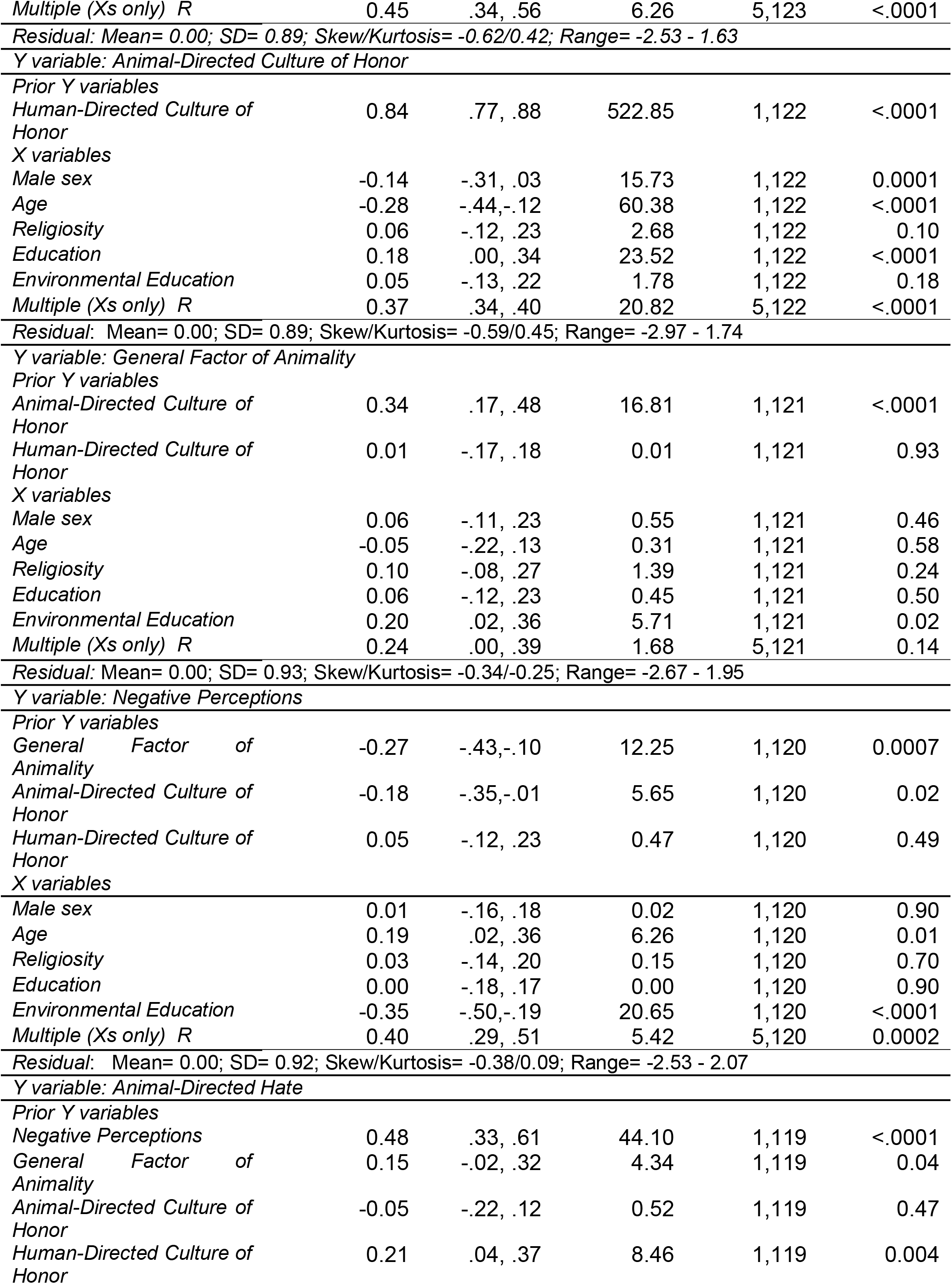
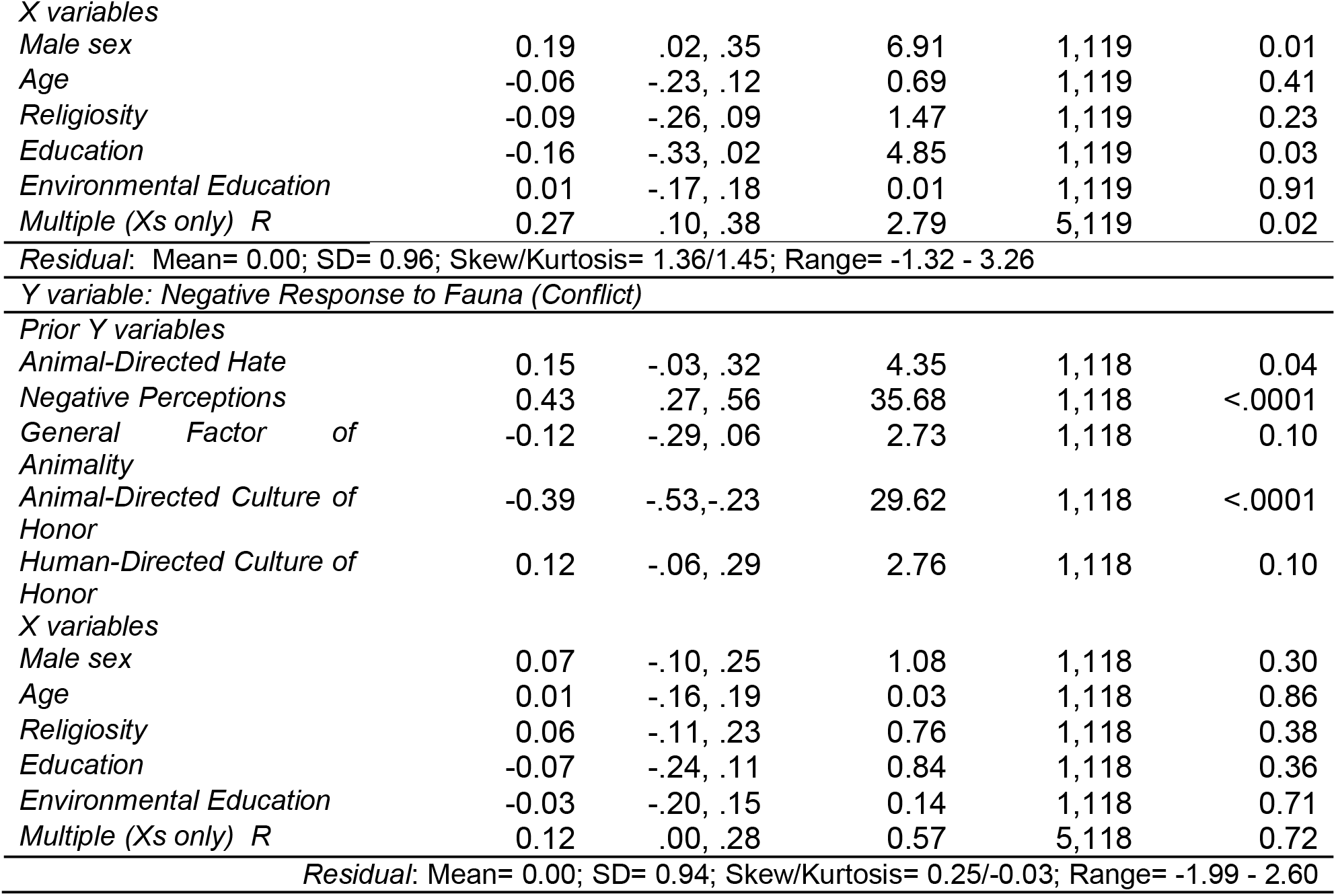
Results of the Sequential Canonical Cascade Model. A Sequential Canonical Cascade model examining the influence of sex, age, religiosity, education, environmental education, human-directed culture of honor, animal-directed culture of honor, a General Factor of Animality, negative perceptions, and animal-directed hate, on participants’ response to fauna.

| *Model* | *Effect size (E)* | *CI* | *F-value* | *df1,df2* | *p-value* |
| --- | --- | --- | --- | --- | --- |
| Overall ($V = 0.651$) | 0.33 | .00,1.00 | 3.04 | 30,610 | <.0001 |
| *Variables* | *Effect size (sR)* | *CI* | *F-value* | *df1,df2* | *p-value* |
| *Y variable: Human-Directed Culture of Honor* |  |  |  |  |  |
| *X variables* |  |  |  |  |  |
| *Male sex* | 0.21 | .03, .37 | 6.66 | 1,123 | 0.01 |
| *Age* | 0.04 | -.14, .21 | 0.25 | 1,123 | 0.62 |
| *Religiosity* | 0.33 | .17, .48 | 17.24 | 1,123 | <.0001 |
| *Education* | 0.21 | .04, .37 | 7.05 | 1,123 | 0.009 |
| *Environmental Education* | -0.03 | -.20, .15 | 0.12 | 1,123 | 0.73 |
| *Multiple (Xs only) R* | 0.45 | .34, .56 | 6.26 | 5,123 | <.0001 |
| *Residual: Mean= 0.00; SD= 0.89; Skew/Kurtosis= -0.62/0.42; Range= -2.53 - 1.63* |  |  |  |  |  |
| *Y variable: Animal-Directed Culture of Honor* |  |  |  |  |  |
| *Prior Y variables* |  |  |  |  |  |
| *Human-Directed Culture of Honor* | 0.84 | .77, .88 | 522.85 | 1,122 | <.0001 |
| *X variables* |  |  |  |  |  |
| *Male sex* | -0.14 | -.31, .03 | 15.73 | 1,122 | 0.0001 |
| *Age* | -0.28 | -.44, -.12 | 60.38 | 1,122 | <.0001 |
| *Religiosity* | 0.06 | -.12, .23 | 2.68 | 1,122 | 0.10 |
| *Education* | 0.18 | .00, .34 | 23.52 | 1,122 | <.0001 |
| *Environmental Education* | 0.05 | -.13, .22 | 1.78 | 1,122 | 0.18 |
| *Multiple (Xs only) R* | 0.37 | .34, .40 | 20.82 | 5,122 | <.0001 |
| *Residual: Mean= 0.00; SD= 0.89; Skew/Kurtosis= -0.59/0.45; Range= -2.97 - 1.74* |  |  |  |  |  |
| *Y variable: General Factor of Animality* |  |  |  |  |  |
| *Prior Y variables* |  |  |  |  |  |
| *Animal-Directed Culture of Honor* | 0.34 | .17, .48 | 16.81 | 1,121 | <.0001 |
| *Human-Directed Culture of Honor* | 0.01 | -.17, .18 | 0.01 | 1,121 | 0.93 |
| *X variables* |  |  |  |  |  |
| *Male sex* | 0.06 | -.11, .23 | 0.55 | 1,121 | 0.46 |
| *Age* | -0.05 | -.22, .13 | 0.31 | 1,121 | 0.58 |
| *Religiosity* | 0.10 | -.08, .27 | 1.39 | 1,121 | 0.24 |
| *Education* | 0.06 | -.12, .23 | 0.45 | 1,121 | 0.50 |
| *Environmental Education* | 0.20 | .02, .36 | 5.71 | 1,121 | 0.02 |
| *Multiple (Xs only) R* | 0.24 | .00, .39 | 1.68 | 5,121 | 0.14 |
| *Residual: Mean= 0.00; SD= 0.93; Skew/Kurtosis= -0.34/-0.25; Range= -2.67 - 1.95* |  |  |  |  |  |
| *Y variable: Negative Perceptions* |  |  |  |  |  |
| *Prior Y variables* |  |  |  |  |  |
| *General Factor of Animality* | -0.27 | -.43, -.10 | 12.25 | 1,120 | 0.0007 |
| *Animal-Directed Culture of Honor* | -0.18 | -.35, -.01 | 5.65 | 1,120 | 0.02 |
| *Human-Directed Culture of Honor* | 0.05 | -.12, .23 | 0.47 | 1,120 | 0.49 |
| *X variables* |  |  |  |  |  |
| *Male sex* | 0.01 | -.16, .18 | 0.02 | 1,120 | 0.90 |
| *Age* | 0.19 | .02, .36 | 6.26 | 1,120 | 0.01 |
| *Religiosity* | 0.03 | -.14, .20 | 0.15 | 1,120 | 0.70 |
| *Education* | 0.00 | -.18, .17 | 0.00 | 1,120 | 0.90 |
| *Environmental Education* | -0.35 | -.50, -.19 | 20.65 | 1,120 | <.0001 |
| *Multiple (Xs only) R* | 0.40 | .29, .51 | 5.42 | 5,120 | 0.0002 |
| *Residual: Mean= 0.00; SD= 0.92; Skew/Kurtosis= -0.38/0.09; Range= -2.53 - 2.07* |  |  |  |  |  |
| *Y variable: Animal-Directed Hate* |  |  |  |  |  |
| *Prior Y variables* |  |  |  |  |  |
| *Negative Perceptions* | 0.48 | .33, .61 | 44.10 | 1,119 | <.0001 |
| *General Factor of Animality* | 0.15 | -.02, .32 | 4.34 | 1,119 | 0.04 |
| *Animal-Directed Culture of Honor* | -0.05 | -.22, .12 | 0.52 | 1,119 | 0.47 |
| *Human-Directed Culture of Honor* | 0.21 | .04, .37 | 8.46 | 1,119 | 0.004 |
| *X variables* |  |  |  |  |  |
| Male sex | 0.19 | .02, .35 | 6.91 | 1,119 | 0.01 |
| Age | -0.06 | -.23, .12 | 0.69 | 1,119 | 0.41 |
| Religiosity | -0.09 | -.26, .09 | 1.47 | 1,119 | 0.23 |
| Education | -0.16 | -.33, .02 | 4.85 | 1,119 | 0.03 |
| Environmental Education | 0.01 | -.17, .18 | 0.01 | 1,119 | 0.91 |
| Multiple (Xs only) R | 0.27 | .10, .38 | 2.79 | 5,119 | 0.02 |
| *Residual: Mean= 0.00; SD= 0.96; Skew/Kurtosis= 1.36/1.45; Range= -1.32 - 3.26* |  |  |  |  |  |
| *Y variable: Negative Response to Fauna (Conflict)* |  |  |  |  |  |
| *Prior Y variables* |  |  |  |  |  |
| Animal-Directed Hate | 0.15 | -.03, .32 | 4.35 | 1,118 | 0.04 |
| Negative Perceptions | 0.43 | .27, .56 | 35.68 | 1,118 | <.0001 |
| General Factor of Animality | -0.12 | -.29, .06 | 2.73 | 1,118 | 0.10 |
| Animal-Directed Culture of Honor | -0.39 | -.53, -.23 | 29.62 | 1,118 | <.0001 |
| Human-Directed Culture of Honor | 0.12 | -.06, .29 | 2.76 | 1,118 | 0.10 |
| *X variables* |  |  |  |  |  |
| Male sex | 0.07 | -.10, .25 | 1.08 | 1,118 | 0.30 |
| Age | 0.01 | -.16, .19 | 0.03 | 1,118 | 0.86 |
| Religiosity | 0.06 | -.11, .23 | 0.76 | 1,118 | 0.38 |
| Education | -0.07 | -.24, .11 | 0.84 | 1,118 | 0.36 |
| Environmental Education | -0.03 | -.20, .15 | 0.14 | 1,118 | 0.71 |
| Multiple (Xs only) R | 0.12 | .00, .28 | 0.57 | 5,118 | 0.72 |
| *Residual: Mean= 0.00; SD= 0.94; Skew/Kurtosis= 0.25/-0.03; Range= -1.99 - 2.60* |  |  |  |  |  |

**Figure 5.**
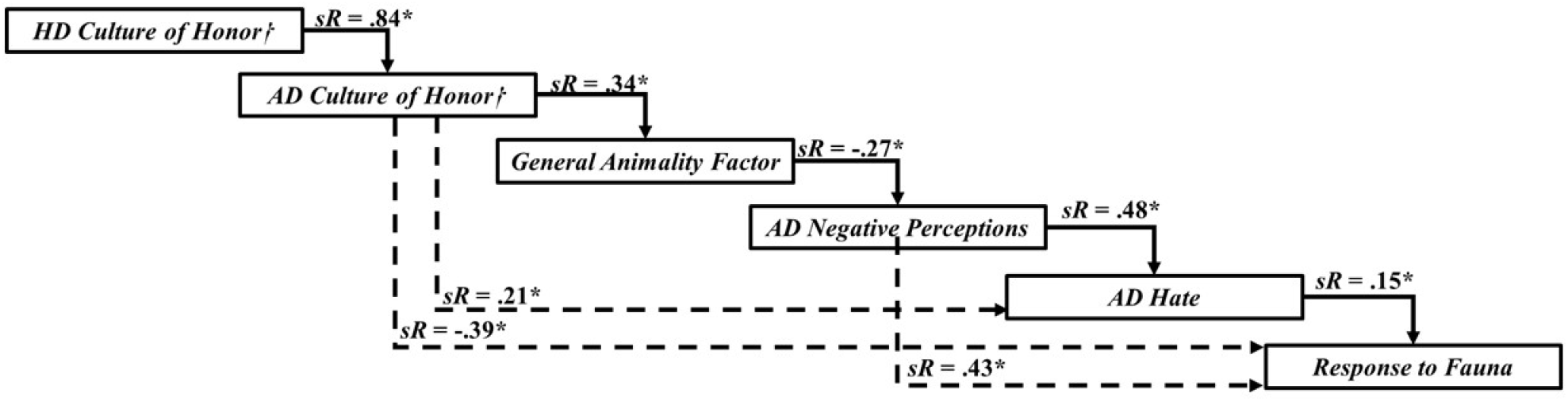
A simplified version of a Sequential Canonical Cascade Model. This Sequential Canonical Cascade Model is examining the influence of human-directed culture of honor, animal-directed culture of honor, a General Factor of Animality, negative perceptions, and animal-directed hate on participants’ response to fauna. Note. The effects of male sex, age, religiosity, education, and environmental education are shown in Table 1. HD: Human-directed; AD: Animal-directed. *p< 0.05; □higher values indicate overreactions.

The animal-directed Culture of Honor scale was positively and sizably influenced by the human-directed Culture of Honor scale. Although education positively predicted the animal-directed Culture of Honor scale (hence, overreaction in the social scenario), this estimate was small in magnitude compared to the effect of the human-directed Culture of Honor scale. Lastly, participants’ age and male gender negatively predicted the animal-directed Culture of Honor scores.

The animal-directed Culture of Honor scores positively predicted the General Factor of Animality. As the omnibus test for the remaining set of predictors did not reach statistical significance, no further interpretation is provided.

The negative perceptions towards animals were negatively predicted by the General Factor of Animality and participants’ environmental education. Participants’ animal-directed hate was positively predicted by the negative perceptions of animals’ factor and the human-directed Culture of Honor scale.

Lastly, participants’ responses to interactions with wild or feral fauna were positively predicted by participants’ animal-directed hate scores, albeit this effect was small in magnitude, and negative perceptions of animals’ factor. Moreover, the animal-directed Culture of Honor scale negatively predicted participants’ responses to conflict. As the omnibus test for the remaining set of predictors did not reach statistical significance, no further interpretation is provided.

## Discussion

Perceptions of wildlife were the strongest predictors of support for retaliatory responses, with hate adding a smaller but consistent effect. An animal-directed Culture of Honor (COH) substantially increased punitive attitudes toward wildlife and was tightly linked to the human-directed honor construct. The General Factor of Animality (GFA) did not directly predict conflict responses but was inversely related to negative perceptions. Environmental education most strongly predicted perceptions (more education generated more favorable views), suggesting an indirect pathway to reducing conflict. Traditional predictors (age, religion) showed limited direct effects once these psychological and cultural variables were modeled.

To our knowledge, most existing literature on HWC and human attitudes toward animals has focused on a limited set of demographic variables, including age, education, and religion, which are often posited to exert significant influence on people’s actions toward animals (Baldry, 2003; Gullone & Robertson, 2008; Chikezie et al., 2023; Ericsson & Heberlein, 2003; Conover, 2002; Pasaribu et al., 2021; White, 1967). However, our findings suggest that human attitudes toward wildlife, particularly in the context of HWC, are more profoundly shaped by a combination of evolutionary and cultural processes, alongside individual- and community-level variables that have historically and continue to influence human–animal interactions.

The construct of Culture of Honor, validated across multiple national contexts, demonstrates that evolutionary and cultural processes shape individuals and communities to exhibit varying propensities for honor-related behavior. Specifically, COH reflects the extent to which individuals respond vengefully to perceived threats to their reputation, property, or social standing, often engaging in extreme violence to avoid appearing weak (Nisbett & Cohen, 1996; Shackelford, 2005). Elevated levels of honor are particularly prevalent in pastoral societies, which are frequently highly dependent on livestock for economic subsistence; in Ecuador, livestock often represents both livelihood and property (Figueredo et al., 2004; Cisneros et al., 2025). Accordingly, we hypothesized that COH might also influence human-wildlife interactions, in addition to intra-human social dynamics. Our results indicate that livestock depredation acts as a trigger for violent retaliatory behaviors consistent with honor-related norms. Moreover, the traditional, human-directed COH construct was strongly correlated with an analogous, animal-directed COH measure, suggesting that individuals who endorse retaliatory aggression in interpersonal contexts are also more likely to exhibit vengeful and aggressive behaviors toward wildlife. This relationship may partially explain the extreme and seemingly disproportionate violence directed at large carnivores (Moscoso, 2020).

Consistent with previous literature, the animal-directed COH also significantly predicted the General Factor of Animality. The construct of Animality is rooted in evolutionary principals, positing that humans’ emotional and cognitive dispositions toward different species – such as empathy, attachment, concern, and curiosity – reflect the historical extent to which those species contributed to human survival and well-being (Keller & Wilson, 1993; Figueredo et al., 2023; Peñaherrera-Aguirre et al., 2024). Consequently, individuals with higher Animality scores, representing a stronger evolutionary affinity and prosocial orientation toward non-human species, are expected to display lower tendencies and support for violent, honor-based retaliation toward wildlife.

Furthermore, the GFA significantly predicted the factor of species perceptions. This finding indicates that contemporary human perceptions of animals are, at least in part, shaped by the evolutionary relationships historically established with them. In the present study, negative perceptions of species emerged as the strongest predictors of HWC responses. Individuals expressing markedly negative views toward the studied species were also those most willing to inflict violence upon or kill those same animals. This pattern helps to elucidate complex global trends in which communities holding unfavorable perceptions of certain species continue to persecute them, even in the absence of actual damage to crops or livestock, and despite their potential ecological or agricultural benefits. Such actions have been widely documented (Reading et al., 2009; Siex & Struhsaker, 1999; Naughton-Treves & Treves, 2005), including in regions where no verified sightings of the targeted species have occurred near agricultural or pastoral areas (Albarracín & Aliaga-Rossel, 2018). Collectively, these results support the view that both the assessed individual perceptions of species and the evolutionary shaping of human– animal bonds - conceptualized through the construct of Animality, which also partially influences present perceptions - are key to understanding the underlying mechanisms driving HWC. This underscores the importance of understanding the origins of human attitudes toward vilified and persecuted species.

Environmental education significantly predicted perceptions of species, indicating that, while evolutionary forces shape views toward wildlife, sustained education can meaningfully influence them. Although it did not directly predict HWC responses, it was the strongest predictor of perceptions, highlighting its potential as a key intervention to reduce species persecution and indirectly mitigate broader conflict outcomes. However, the environmental education measure captured cumulative exposure over individuals’ lifetimes and academic trajectories. These results should not be taken to imply that brief or one-time environmental education sessions are sufficient; rather, they highlight the long-term benefits of sustained, early exposure in fostering awareness, tolerance, and reducing the likelihood of HWC (Ericsson & Heberlein, 2003; Conover, 2002).

Finally, hate scores were the only remaining variable showing a modest correlation with responses to HWC. Despite the small magnitude of this correlation, there is a clear association between hate scores and the perception scale, consistent with prior literature demonstrating strong links between emotions and perceptions of animals (Prokop, Fancovicova, & Kubiatko, 2009). These findings suggest that individuals with higher hate scores are more likely to hold negative perceptions of species and, consequently, may be more prone to engage in violent responses toward wildlife.

Among all variables included in the model, negative responses to conflict scenarios involving wildlife were most strongly predicted by negative perceptions, followed by the animal-directed COH and to a lesser extent by individuals’ hate toward species. These findings may provide a valuable tool for diagnosing and assessing the severity of HWC and the degree of vulnerability faced by wild fauna across communities and regions. High levels of negative perceptions and honor within a community substantially increase the risk that wildlife species will be subjected to poaching, persecution, violent retaliation, and eradication, even when the species poses little or no actual harm. Applying these tools to identify areas of high vulnerability can therefore play a critical role in the conservation of endangered and highly persecuted species, such as the spectacled bear (*Tremarctos ornatus*) in the Ecuadorian Andes (MAE, 2019; Vela-Vargas et al., 2021; García-Rangel, 2012; Cisneros et al., 2025).

### Future directions

The present study suggests that evolutionary forces strongly shape human attitudes toward different species. Applying these methodological tools across regions could enable cross-cultural comparisons, providing insight into the distinct evolutionary trajectories communities have experienced with specific species. Such comparisons could also reveal species’ relative vulnerability, informing conservation priorities and highlighting areas at greatest risk of extirpation. The constructs of Culture of Honor, Animality, and perceptions offer accessible diagnostic measures for assessing these risks, which, when integrated into regional HWC risk maps, could help identify vulnerable hotspots.

### Limitations

A limitation of this study is the challenge of recruiting large samples, given that the study area comprises small peri-rural communities in southern Ecuador adjacent to forested wildlife habitats. Nevertheless, relative to the overall population size, the obtained sample is sufficient to provide meaningful insights. Future research should aim to include a broader range of communities to increase sample sizes and enhance the generalizability of findings.

## Conclusions

Our study demonstrates that evolutionary and cultural variables exert a significant and previously underappreciated influence on HWC responses, surpassing the explanatory power of commonly studied sociodemographic factors. These findings underscore the practical value of incorporating evolutionary perspectives into conflict mitigation, as they can serve diagnostically to identify regions where retaliatory actions against wildlife are most likely, thereby allowing the prioritization of interventions and the effective allocation of resources. Implementing these tools across conflict-prone landscapes represents a concrete next step toward reducing HWC and conserving endangered carnivores, while also advancing our understanding of the origins of human attitudes toward animals.

## Supporting information

Interview in English

## Notes

### Competing Interest Statement

The authors have declared no competing interest.

