## Supplementary material for "The role and origins of human attitudes in Human–Wildlife Conflict responses: Insights from Spectacled Bears (*Tremarctos ornatus*) and other wild carnivores in Southern Ecuador": Interview in English

### Interview on Human-Wildlife Conflicts in Strategic Areas

**Interview Code** (Locality Code / Interviewer Initials / Interview Number) ___/___/___/

**General Survey Information:**

Interviewer Name: ________________________________________ Day: ______ Month: ______ Year: 20____
Start Time: _________________ Coordinates: N ______________________ E__________________________
Locality: __________________________ Parish: ____________________ Canton: _______________________

##### A) General Information

1) Gender: _____ 2) Age (in completed years): _____ 3) Place of Birth: _________________________
4) Place of Residence: _______________ 5) Are you a resident of the area? _____ (If "No," skip to Question 7)
6) How long have you lived in the area? (years or months) ______________ 7) How long have you worked in the area? _________________ 8) Number of family members: _______________________

##### B) Education

9) What is the highest level of education you have completed? ____________________________
(A) No education (skip to Question 10), (B) Literacy center, (C) Primary, (D) High school, (E) Higher education, (F) Postgraduate, (G) Other (please specify): ____________________________
10) Are you currently attending an educational institution? Yes/No_____
11) What is the highest level of education your father has completed? _______________________
(A) No education (skip to Question 12), (B) Literacy center, (C) Primary, (D) High school, (E) Higher education, (F) Postgraduate, (G) Other (please specify): ____________________________
12) What is the highest level of education your mother has completed? _____________________
(A) No education (skip to Question 13), (B) Literacy center, (C) Primary, (D) High school, (E) Higher education, (F) Postgraduate, (G) Other (please specify): ____________________________

##### C) Environmental Education

*Please indicate how often the following statements apply to you. Use the scale below and note your responses in the spaces provided.*

| 1 | 2 | 3 | 4 | 5 |
|---|---|---|---|---|
| Never | Rarely | Sometimes | Often | Very often |

13) How frequently, during your life, have someone spoken with you about taking care of the environment. __________________________
14) How frequently, during your life, have they spoken with you about the importance of nature. ____________________________
15) How frequently, during your life, have they spoken with you about taking care of the forest. ____________________________
16) How frequently, during your life, have they spoken with you about the importance of wildlife. ___________________________
17) How frequently, in school, have they spoken with you about taking care of the environment. __________________________

18) How frequently, in school, have they spoken with you about the importance of nature.

______________________________

19) How frequently, in school, have they spoken with you about taking care of the forest.

______________________________

20) How frequently, in school, have they spoken with you about the importance of wildlife.

_____________________________

**D) Religion**

21) The centrality of Religiosity Scale (CRS) (2012 Stefan Huber & Odilo W. Huber)

*Please indicate how often the following statements apply to you. Use the scale below and note your responses in the spaces provided. Frequency scale is used from on questions 1, 3, 4, 5 and 10, importance scale on 2, 6, 7, 8, 9.*

| **Never**<br><br>**Not at all** | **Rarely**<br><br>**Not very much** | **A few times a year**<br><br>**Moderately** | **One or three times a month**<br><br>**Quite a bit** | **Once or multiple times a week**<br><br>**Very much so** |
|---|---|---|---|---|
| **1** | **2** | **3** | **4** | **5** |

1. How often do you think about religious issues? _______

2. To what extent do you believe that God or something divine exists? _______

3. How often do you take part in religious services? _______

4. How often do you pray? _______

5. How often do you experience situations in which you have the feeling that God or something divine intervenes in your life? _______

6. How interested are you in learning more about religious topics? _______

7. To what extend do you believe in an afterlife—e.g. immortality of the soul, resurrection of the dead or reincarnation? _______

8. How important is to take part in religious services? _______

9. How important is personal prayer for you? _______

10. How often do you experience situations in which you have the feeling that God or something divine wants to communicate or to reveal something to you? _______

**E) Animality:**

*Please indicate how strongly you agree or disagree with the following statements. Use the scale below and note your responses in the spaces provided.*

| Strongly Disagree | Slightly Disagree | Neither agree nor disagree | Slightly Agree | Strongly Agree |
|---|---|---|---|---|
| 1 | 2 | 3 | 4 | 5 |

1. I really love animals. _______

2. I feel upset when I see animals in pain. _______

3. I don't believe that people should treat animals like family members. _______

4. I enjoy learning about animals. _______

5. Sometimes I sense a special energy between me and animals. _______

6. Animals mean more to me than my friends. _______

7. I have a loving relationship with animals. _______

8. I am humane and soft-hearted towards animals. _______

9. I feel that animals should always be kept outside. _______

10. I have a lot of intellectual curiosity about animals. _______

11. Sometimes I feel that I have a gift or a sixth sense for understanding animals. _______

12. I prefer to be around animals more than people. _______

13. I like animals because they bring me joy and unconditional friendship. _______

14. I am filled with joy when I see animals playing and frolicking. _______

15. I think some animals should be considered as part of a family. _______

16. I am strongly interested in the behavior of animals. _______

17. I enjoy the special connection I have with animals. _______

18. I have a better relationship with animals than with people. _______

19. I am comfortable around animals. _______

20. If necessary, it's ok for people to harm animals to serve the needs of humans. _______

21. I notice more details about animals and their behavior than most people. _______

22. I am caring and affectionate with animals. _______

23. I am somewhat tense or uneasy when around animals. _______

24. If necessary, I am willing to manipulate or use animals to serve my needs. _______

25. I have a good intuitive sense about animals. _______

26. I think animals deserve our affection. _______

27. Usually, I feel at ease around animals. _______

28. I don't think animals should ever be used as food for people. _______

29. I have a good understanding of what animals are feeling. _______

30. When I see an animal, I often have the impulse to want to pet it or hold it. _______

**F) Culture of Honor:**

*Please indicate how strongly you agree or disagree with the following statements. Use the scale below and note your responses in the spaces provided.*

| 1 | 2 | 3 | 4 | 5 |
|---|---|---|---|---|
| Did much less than was justified | Did somewhat less than was justified | Did neither more nor less than was justified | Did somewhat more than was justified | Did much more than was justified |

1. A man punches a drunk who bumped into the man's wife on the street. _______
2. A woman shoots another person because that person had sexually assaulted the woman's sister. _______
3. A man seduces the daughter of another person because that person had seduced his 16-year-old daughter. _______
4. A woman stabs an adult male who had previously beaten up her mother. _______
5. A man fights a friend when they are having an argument because his friend called him a liar and a coward to his face. _______
6. A woman insults another person because that person calls her a liar and a cheat. _______
7. A man fights an acquaintance for looking over the man's girlfriend and talking to her in a suggestive way, even if she was not offended by it. _______
8. A woman fights another woman who deeply insulted her in public. _______
9. A man hits a woman back for slapping him in the face when he gets fresh with her. _______
10. A woman reveals a female friend's secrets because that friend has offended her. _______
11. A man fights an acquaintance because that acquaintance looks over the man's girlfriend and starts talking to her in an offensive way. _______
12. A man punches an adult stranger who was in a protest march showing opposition to the man's views. _______
13. A woman hits a man who stole a statue of the Virgin Mary from her church. _______

**G) Culture of Honor (adapted to animals):**

*Please indicate how strongly you agree or disagree with the following statements. Use the scale below and note your responses in the spaces provided.*

| 1 | 2 | 3 | 4 | 5 |
|---|---|---|---|---|
| Did much less than was justified | Did somewhat less than was justified | Did neither more nor less than was justified | Did somewhat more than was justified | Did much more than was justified |

1. A man throws a stone at a forest animal that was trying to attack his calf. _______
2. A woman shoots a forest animal because it had attacked her cow. _______
3. A man tries to steal the offspring of a forest animal that had been circling the pasture where his cows were. _______
4. A woman slashes a forest animal with a machete after it had previously attacked her best cow. _______
5. A man throws a stone at a forest animal that had repeatedly stolen his chickens, and he was the only one in the community experiencing this. _______
6. A woman yells at a forest animal that had repeatedly damaged her crops, and she was the only one in the community affected by this. _______
7. A man throws a stone at a forest animal that had been trying to enter his chicken coop. _______
8. The community had captured and tied up a forest animal in the town square. A man kicks it in the face because it scared him with a roar. _______
9. The community had captured and tied up a forest animal in the town square. A woman kicks it in the face because the animal tried to catch her. _______
10. A woman tells her neighbors where the nest of a forest bird is, as it had attempted to steal her chickens. _______
11. A man was friends with a forest animal that had lived near his farm for years. One day, the man threw a stone at the animal because it attacked his chickens. _______
12. A forest animal lived near a farm. It had never caused any harm, but the day it entered the farm, the farmer threw a stone at its head. _______
13. A woman throws a stone at a forest bird that entered the town church and knocked down the statue of the Virgin Mary. ______

**H) Reactions Regarding Wildlife**

22) What would your reaction be if you encountered the following wildlife?

| Reaction/ Species | Species 1 | Species 2 | Species 3 | Species 4 | Species 5 | Species 6 |
|---|---|---|---|---|---|---|
| Flee the area | | | | | | |
| Attack the animal | | | | | | |
| Scare the animal | | | | | | |
| Capture the animal | | | | | | |
| Hunt the animal | | | | | | |
| Call the authorities | | | | | | |
| Do nothing | | | | | | |
| Other (please specify) | | | | | | |

**I) Emotions towards wildlife:**

Each species must be ranked from 1 to 5 on how the interviewee feels about it. For example: how much does the interviewee fear the bear from 1 to 5, and so on.

| | Species 1 | Species 2 | Species 3 | Species 4 | Species 5 | Species 6 |
|---|---|---|---|---|---|---|
| Hatred | | | | | | |

**J) Perceptions towards wildlife:**

Each species must be ranked from 1 to 5 on how the interviewee feels about it. For example: how important does the interviewee think the bear is from 1 to 5, and so on.

| | Species 1 | Species 2 | Species 3 | Species 4 | Species 5 | Species 6 |
|---|---|---|---|---|---|---|
| Dangerous | | | | | | |
| Aggressive | | | | | | |
| Destructive | | | | | | |
| Unpredictable | | | | | | |
| Greedy | | | | | | |
| Threat to Livestock | | | | | | |
| Economic loss (how much) | | | | | | |
| Damage (how much) | | | | | | |
| Cute | | | | | | |
| Important for my culture | | | | | | |
| Important to nature | | | | | | |
| Peaceful | | | | | | |
| Impressive | | | | | | |
| Playful | | | | | | |
| Inspiring | | | | | | |

End Time: _____________

**Editing of the Interview Structure:** F. Lopes, 2024. **Reference Materials Used:** R. Cisneros, 2018.
